# Enhancing TFEA.ChIP with ENCODE Regulatory Maps for Generalizable Transcription Factor Enrichment

**DOI:** 10.1101/2025.07.28.667192

**Authors:** Yosra Berrouayel, Luis del Peso

**Affiliations:** Instituto de Investigaciones Biomédicas Sols-Morreale (IIBM) CSIC-UAM, Departamento de Bioquímica, Universidad Autónoma de Madrid (UAM), Arturo Duperier, 4, 28029, Madrid, spain; IdiPaz, Instituto de Investigación Sanitaria del Hospital Universitario La Paz, Pedro Rico, 6, 28029, Madrid, Spain; Centro de Investigación Biomédica en Red de Enfermedades Respiratorias (CIBERES), Instituto de Salud Carlos III, Melchor Fernández Almagro, 3, 28029, Madrid, Spain; Unidad Asociada de Biomedicina, Consejo Superior de Investigaciones Científicas, Universidad de Castilla - La Mancha, 02006, Albacete, Spain

**Keywords:** gene regulation, TF enrichment, enhancer, Regulatory Elements to Genes

## Abstract

Identifying transcription factors (TFs) responsible for gene expression changes remains a central challenge in functional genomics. TFEA.ChIP is a ChIP-seq–based TF enrichment analysis tool that addresses this by linking TF binding profiles to differentially expressed genes through experimentally supported cis-regulatory element (CRE)–gene associations. Unlike motif- or heuristic-based approaches, TFEA.ChIP adopts a biologically grounded strategy by intersecting TF binding data from ReMap2022 with regulatory maps from ENCODE’s rE2G and CREdb. To overcome the high context-specificity of rE2G associations, we developed filtering strategies based on confidence scores and recurrence across biosamples. Benchmarking on 369 curated gene sets from the MSigDB C2 CGP collection showed that recurrence-based filtering significantly improved accuracy, outperforming the original GeneHancer-based implementation and leading tools including BARTv2.0, Lisa, ChEA3, and HOMER. A case study on hypoxia further validated the method, demonstrating accurate and pathway-specific enrichment of HIF-related TFs using both overrepresentation analysis and gene set enrichment analysis (GSEA). Additionally, the updated implementation of TFEA.ChIP in R/Bioconductor introduces several user-friendly features, including automated analysis workflows and expression-based filtering of candidate TFs. These additions streamline the integration of TFEA.ChIP into standard RNA-seq analysis pipelines, enabling more efficient and reproducible workflows. Together with its strong benchmarking performance and biologically grounded framework, the updated tool provides a robust and accessible solution for inferring transcriptional regulators from gene expression data.

## Introduction

One of the central challenges in transcriptional biology is deciphering which transcription factors (TF) are responsible for the observed changes in gene expression. While gene expression profiling returns lists of differentially expressed genes, identifying the upstream regulators driving these changes remains non-trivial. To address this, we present an updated version of TFEA.ChIP, a TF enrichment analysis tool that leverages ChIP-seq datasets to identify TFs most likely regulating a given gene set.

Several tools exist for inferring regulatory TFs from gene lists, including BARTv2.0 [12], Lisa [15], ChEA3 [9] and HOMER [7]. These tools use different methodologies: BARTv2.0 infers TF regulators by comparing the query gene set to a large compendium of ChIP-seq profiles across multiple cell types using an enrichment score based on epigenomic similarity; Lisa combines DNase-seq and ChIP-seq data with a machine learning model to predict the TFs most likely to regulate a given gene set; ChEA3 integrates TF–target associations from diverse sources, including ChIP-seq data, co-expression analyses, and literature mining, to rank TFs based on multiple gene set libraries; HOMER focuses primarily on motif enrichment within regulatory regions, using predefined TF binding motifs to identify overrepresented sequences. In contrast, TFEA.ChIP adopts a direct and interpretable approach: it defines the target genes of each TF based on experimentally derived ChIP-seq binding peaks from uniformly processed datasets from ReMap2022 [5], and assesses enrichment using either Fisher’s exact test for an overrepresentation analysis (ORA), or Gene Set Enrichment Analysis (GSEA) (Figure 1). This method prioritizes transparency and biological grounding over model complexity.

**Fig. 1.**
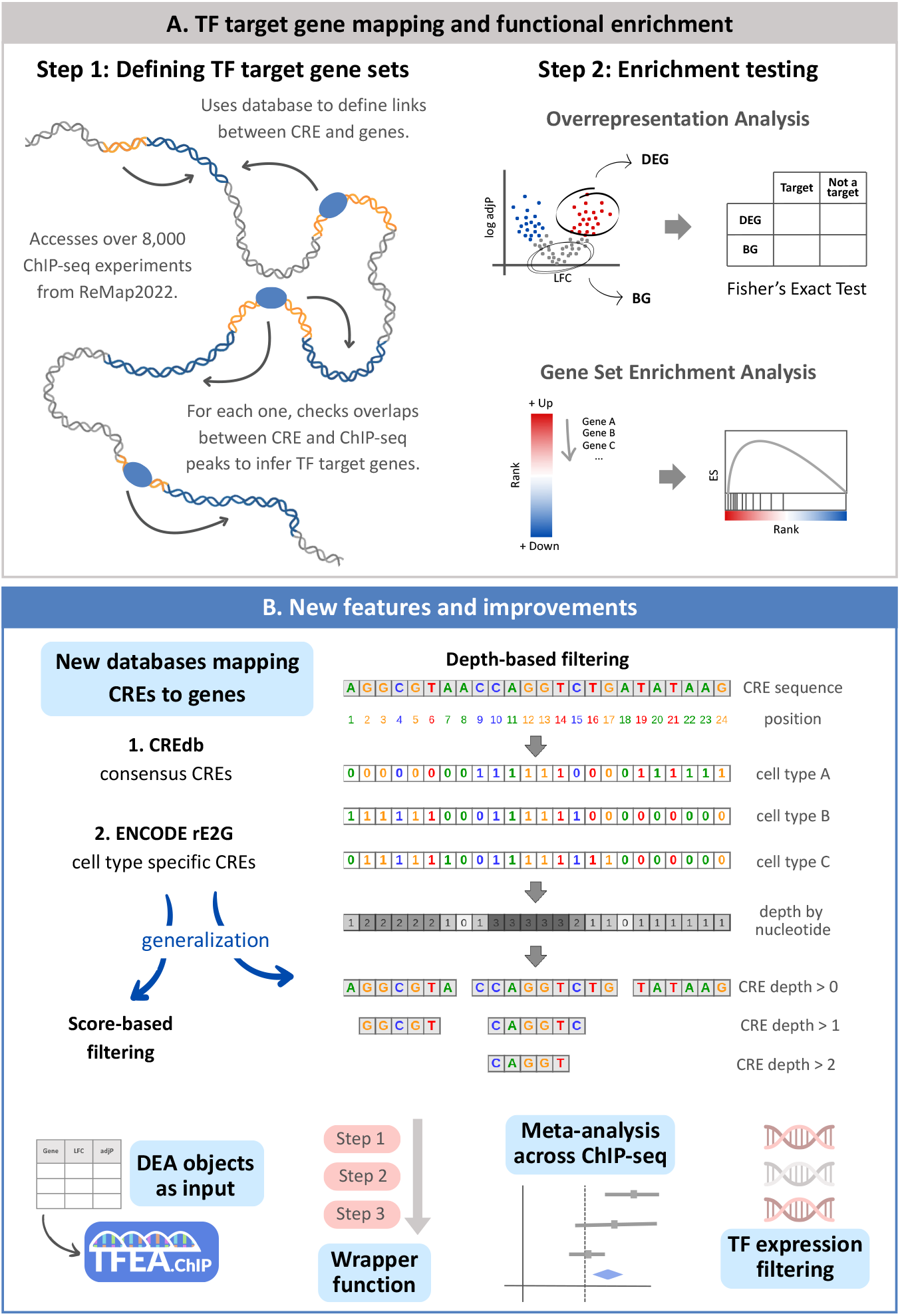
Overview of the TFEA.ChIP framework and methodological enhancements. The A panel illustrates the two-step core workflow of TFEA.ChIP. In step 1, TF target gene sets are defined using external databases that map CREs to their putative target genes. ChIP-seq peaks from ReMap2022 are then intersected with these CREs to identify the corresponding TF targets. In step 2, enrichment analysis is conducted to evaluate whether the identified TF target genes are significantly associated with a user-provided gene list. Two complementary approaches are supported: Overrepresentation Analysis, which uses Fisher’s exact test to assess enrichment among differentially expressed genes, and Gene Set Enrichment Analysis, which analyzes ranked gene lists to detect shifts in the distribution of TF targets. The B panel highlights key enhancements introduced in the current version of TFEA.ChIP. The tool integrates two new sources for CRE–gene mapping: (1) CREdb, a database of consensus regulatory elements, and (2) ENCODE rE2G, which provides cell-type-specific CRE–gene associations derived from functional genomics data. To extend the applicability of rE2G to cell types not directly profiled, we implemented a score- and depth-based filtering strategy. In this framework, the score quantifies the strength of evidence supporting a CRE–gene association, while depth reflects the number of cell types in which each nucleotide within a CRE is functionally annotated. Additional functionality includes support for direct input of differential expression analysis result objects, a wrapper function that automates the complete enrichment analysis pipeline, a meta-analysis function for aggregating results across multiple ChIP-seq datasets for the same TF, and a TF expression filtering step to exclude non-expressed TFs.

A critical step in TF enrichment analysis is linking cis-regulatory elements (CREs) to their target genes. Tools such as HOMER typically assign CREs to the nearest gene, which may not always reflect true regulatory relationships [16, 1]. TFEA.ChIP, however, bases these associations on experimental evidence. The previous version relied on GeneHancer [3], an integrative resource that connects genes to regulatory elements by aggregating data from ENCODE, FANTOM5, eQTLs, and chromatin interaction datasets. While the previous implementation of TFEA.ChIP achieves strong performance, benefiting from the high-quality enhancer–gene mappings offered by GeneHancer, further enhancements are still possible[11]. To further improve the accuracy of TF enrichment analysis, the updated version of TFEA.ChIP incorporates two additional CRE–gene association resources: rE2G (Regulatory Elements to Genes) developed by the ENCODE Consortium [4], and CREdb [6]. rE2G uses a supervised model trained on large-scale CRISPR perturbation datasets, learning enhancer-gene links using over 13 molecular features such as chromatin accessibility, 3D genome architecture, and enhancer synergy. It provides probabilistic predictions across 352 biosamples, offering tissue-specific regulatory maps validated against eQTL and GWAS datasets. CREdb, on the other hand, provides a curated database of high-confidence CREs, including enhancers, promoters, silencers, and insulators, aggregated from over 11 databases and harmonized across more than 1,000 cell types and tissues. By incorporating these robust and biologically grounded resources, the new version of TFEA.ChIP significantly improves the accuracy of enhancer-gene associations, thereby enhancing the identification of relevant regulatory TFs.

While there is currently no universally accepted gold-standard benchmark for evaluating TF enrichment tools, the C2 CGP collection from the Molecular Signatures Database (MSigDB) [10, 18] serves as a practical and informative proxy. This collection comprises manually curated gene sets derived from a variety of chemical and genetic perturbation experiments, including many where specific TFs have been directly manipulated. As such, it provides a biologically relevant context for testing the ability of enrichment tools to recover known TF-gene associations. Using the C2 CGP gene sets for benchmarking, we compared the updated version of TFEA.ChIP against both its previous release and other state-of-the-art tools. The results demonstrate that the new implementation consistently improves the precision and accuracy of TF identification. This highlights the value of the newly integrated enhancer-gene resources and the robustness of TFEA.ChIP’s enrichment framework.

In summary, the newly integrated resources offer a complementary improvement over GeneHancer by contributing more accurate and experimentally supported CRE-gene associations. This refined regulatory mapping enhances TFEA.ChIP’s ability to identify likely transcriptional regulators. Benchmarking against the previous version and other established tools shows consistent performance gains, underscoring the utility of TFEA.ChIP as a biologically informed approach to transcriptional regulatory analysis.

## Methods

### Definition of databases

We generated multiple versions of the underlying database on which TFEA.ChIP depends, each based on different regulatory maps derived from either ENCODE’s rE2G predictive models or the CREdb database. For the rE2G-based versions, we aimed to construct generalized regulatory maps by aggregating predicted CRE–gene associations across all available biosamples. These links were filtered using two different strategies:

1. Confidence score thresholding. Each CRE–gene pair in rE2G is assigned a probability score derived from a logistic regression model, indicating the likelihood of a true regulatory relationship. We generated three filtered versions by applying score thresholds of 0.25, 0.50, and 0.75, retaining 85.85% (16.6M), 51.87% (10.0M), and 35.72% (6.9M) of links, respectively, compared to the unfiltered 19.3M total. These thresholds were chosen to explore the trade-off between sensitivity and confidence, providing flexible options depending on the analysis goals.

2. Recurrent link filtering (“depth”). In parallel, we implemented a second filtering strategy based on link recurrence across cell types, or “depth”, defined as the number of biosamples in which a CRE–gene pair is predicted (Figure 1). This method prioritizes regulatory links that are consistently observed across diverse biological contexts. Using depth thresholds of *>*50, *>*100, *>*200, and *>*300, we produced increasingly stringent subsets of the rE2G dataset, retaining 64,563 (0.33%), 35,574 (0.18%), 19,135 (0.10%), and 12,546 (0.06%) links, respectively.

CREdb annotations were used without additional filtering. This database integrates over 17 million CRE–gene links curated from 11 public sources, harmonized across more than 1,000 cell types. While CREdb lacks recurrence metrics, it provides breadth and annotation diversity that complements the ENCODE-based rE2G maps.

In addition to the rE2G and CREdb CREs, we included gene promoters, defined as the regions spanning 1 kb upstream and downstream of the transcription start site. Each version of the TFEA.ChIP database was generated using the `makeChIPGeneDB` function included in the TFEA.ChIP R package, incorporating the ReMap2022 ChIP-seq binding datasets [5] with default parameters.

### TFEA.ChIP version benchmarking

#### Overrepresentation analysis

To benchmark the performance of each TFEA.ChIP database version, we conducted an ORA using a curated subset of TF-regulated gene sets from the MSigDB C2 CGP collection. From an initial pool of 3,494 gene sets, we retained 369 sets that met two criteria: (i) the presence of a well-defined upstream regulator and (ii) the availability of at least one corresponding ChIP-seq dataset for the TF in the ReMap2022 database (Supplementary Table 1). Although all gene sets were represented using human gene symbols, some originated from non-human model organisms, predominantly mouse. In these cases, the corresponding human orthologs were identified and used for analysis.

TF enrichment analysis was performed in TFEA.ChIP using the `contingency_matrix` function, which constructs contingency tables comparing ChIP-seq target genes with input gene sets, and the `getCMstats` function, which calculates odds ratios (ORs) from these tables to quantify the strength of association between TF binding and presence in the gene sets. Because MSigDB lacks a standardized background gene list, we conducted the analysis 100 times using different sets of 1,000 randomly selected genes. For each dataset, the mean OR from these randomized runs was used as a robust estimate of enrichment strength for the expected TF. This ORA pipeline was applied consistently across all TFEA.ChIP database versions, including the original GeneHancer-based implementation.

In a case study focused on hypoxia signaling, we applied TFEA.ChIP to a consensus gene expression signature derived from a recent meta-analysis of hypoxia datasets [14]. Differentially expressed genes (DEGs) were defined using a threshold of absolute log_2_ fold change (|LFC|) greater than 0.5 and an adjusted p-value less than 0.05. As background, we used genes with |LFC| *<* 0.5 and adjusted p-value *>* 0.5. We specifically assessed enrichment for HIF complex components (HIF1A, EPAS1, and ARNT) and reported only statistically significant results (p *<* 0.05).

To evaluate the biological relevance of each ChIP-seq dataset, we manually annotated experimental metadata for hypoxia pathway activation. Datasets were classified as “Active” if they were performed under hypoxic conditions, involved treatments known to activate HIF signaling, or used cell types with constitutive HIF pathway activity. Datasets from untreated or normoxic conditions were labeled “Inactive”, and those with unclear or incomplete metadata were classified as “NA” (Not Assigned).

#### Gene Set Enrichment Analysis

To complement the ORA, we performed GSEA using the hypoxia meta-analysis case study. Genes were ranked by their LFC, from the most upregulated to the most downregulated, based on the meta-analysis results. GSEA was carried out using the `GSEA_run` function provided in the TFEA.ChIP package. This analysis was repeated independently for each version of the regulatory database to compare the enrichment results across different CRE-gene mapping strategies. Only results with nominal p-values *<* 0.05 were considered statistically significant and retained for interpretation. Then, as in the ORA, we focused on effect sizes, in this case the enrichment scores (ES) obtained for the core hypoxia-inducible TFs: HIF1A, EPAS1, and ARNT. To contextualize the biological relevance of each ChIP-seq dataset, we applied the same hypoxia pathway activity annotations described above. Datasets were classified as “Active” if generated under hypoxic conditions, “Inactive” if collected under normoxic or untreated conditions, and “NA” (Not Assigned) when the metadata was ambiguous or insufficient to determine pathway status.

### Performance evaluation using AUC

To quantitatively assess the predictive performance of each TFEA.ChIP database version, we computed the Area Under the Receiver Operating Characteristic Curve (AUC) using the `pROC` package in R. For each gene set included in the selected C2 CGP collection from MSigDB, the ChIP-seq datasets corresponding to the TF known to regulate that gene set were designated as the “positive class”. To construct the “negative class”, an equal number of ChIP-seq datasets were randomly selected from TFs not known to regulate the gene set. This design was used to address the substantial class imbalance, as the number of ChIP-seq datasets from true regulatory TFs is typically much smaller than the number from non-regulatory TFs.

AUC computation was repeated 100 times per gene set, with a different random sampling of negative TFs in each iteration. In every iteration, all ChIP-seq datasets were assigned a score based on their rank in the enrichment output using the following formula:

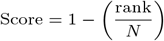

where *rank* denotes the position of the ChIP-seq dataset in the sorted list (with lower ranks indicating stronger enrichment), and *N* is the total number of ChIP-seq datasets considered.

For each gene set, the final AUC score was calculated as the median across the 100 iterations, providing a robust and stable estimate of the tool’s ability to prioritize true regulators over background noise.

## Results

### Definition of CRE–gene databases and comparative evaluation

Establishing reliable links between CREs and their target genes remains a critical and complex step in identifying TF targets, particularly for enhancers acting at a distance from promoters. These enhancer–gene associations are often highly cell-type specific [8], which poses challenges for tools like TFEA.ChIP that aim to be broadly applicable across diverse biological contexts. To address this, we constructed a unified CRE–gene database that integrates publicly available resources (Figure 1A) while balancing breadth of coverage with reliability.

As a primary resource, we incorporated all CRE–gene links predicted by the rE2G model, a supervised machine learning framework trained on functional genomic datasets and CRISPR perturbation experiments to infer CRE–gene associations across 352 biosamples. While this comprehensive integration maximizes coverage, it also introduces potential noise due to the inclusion of highly context-specific associations. To mitigate this, we implemented two filtering strategies designed to enhance both confidence and generalizability of the CRE–gene links (Figure 1B). First, we applied score-based filtering using the rE2G confidence score, a metric that reflects the strength of evidence supporting each enhancer–gene pair, based on molecular features. We defined three filtered subsets with minimum confidence thresholds of 0.25, 0.50 and 0.75, denoted rE2G25s, rE2G50s, and rE2G75s, respectively. The complete, unfiltered dataset is referred to as rE2G. Second, we introduced a depth-based filtering approach, where “depth” refers to the number of biosamples in which a given enhancer–gene association is observed. This captures the recurrence of regulatory links across diverse cellular contexts. We constructed four filtered datasets with depth thresholds of *>*50, *>*100, *>*200, and *>*300 biosamples, designated rE2G50d, rE2G100d, rE2G200d, and rE2G300d.

To evaluate the impact of these filtering strategies on TF enrichment performance, we selected a benchmark of 369 gene sets from the MSigDB C2 CGP collection [10, 18]. These gene sets were selected for their association with chemical or genetic perturbations in which the regulating TF is explicitly annotated and has corresponding ChIP-seq data in ReMap2022 (see *Methods*). Each gene set was analyzed with TFEA.ChIP using the different versions of the databases described above and applying ORA, which calculates both log_2_OR and adjusted p-values to assess the enrichment of TF binding within the gene set (Figure S1). To assess and compare performance across database versions, we extracted the log_2_ OR from all TF ChIP-seq profiles associated with the benchmarking gene sets (Figure 2A). These results indicate that the unfiltered rE2G dataset underperformed relative to the original TFEA.ChIP implementation, producing lower log_2_OR values for the relevant TFs in the vast majority of cases. Applying score-based filtering to the rE2G data progressively increased enrichment performance, with higher confidence thresholds resulting in stronger enrichment signals. Depth-based filtering, however, had a notably stronger and more consistent impact. Increasing the minimum depth threshold resulted in more pronounced enrichment signals, with performance consistently surpassing that of both the original method and the score-filtered versions. This was evidenced by higher log_2_OR values for the relevant TFs in nearly all tested cases. This conclusion is further supported by AUC analysis (Figure 2B), which evaluates each method’s ability to rank relevant TFs above non-relevant ones. Among all tested configurations, the rE2G300d version achieved the highest AUC values, slightly outperforming alternative filtering strategies and all baseline versions. Consequently, it represents the preferred database for CRE–gene mapping and it is the one used by default in the new version of TFEA.ChIP.

**Fig. 2.**
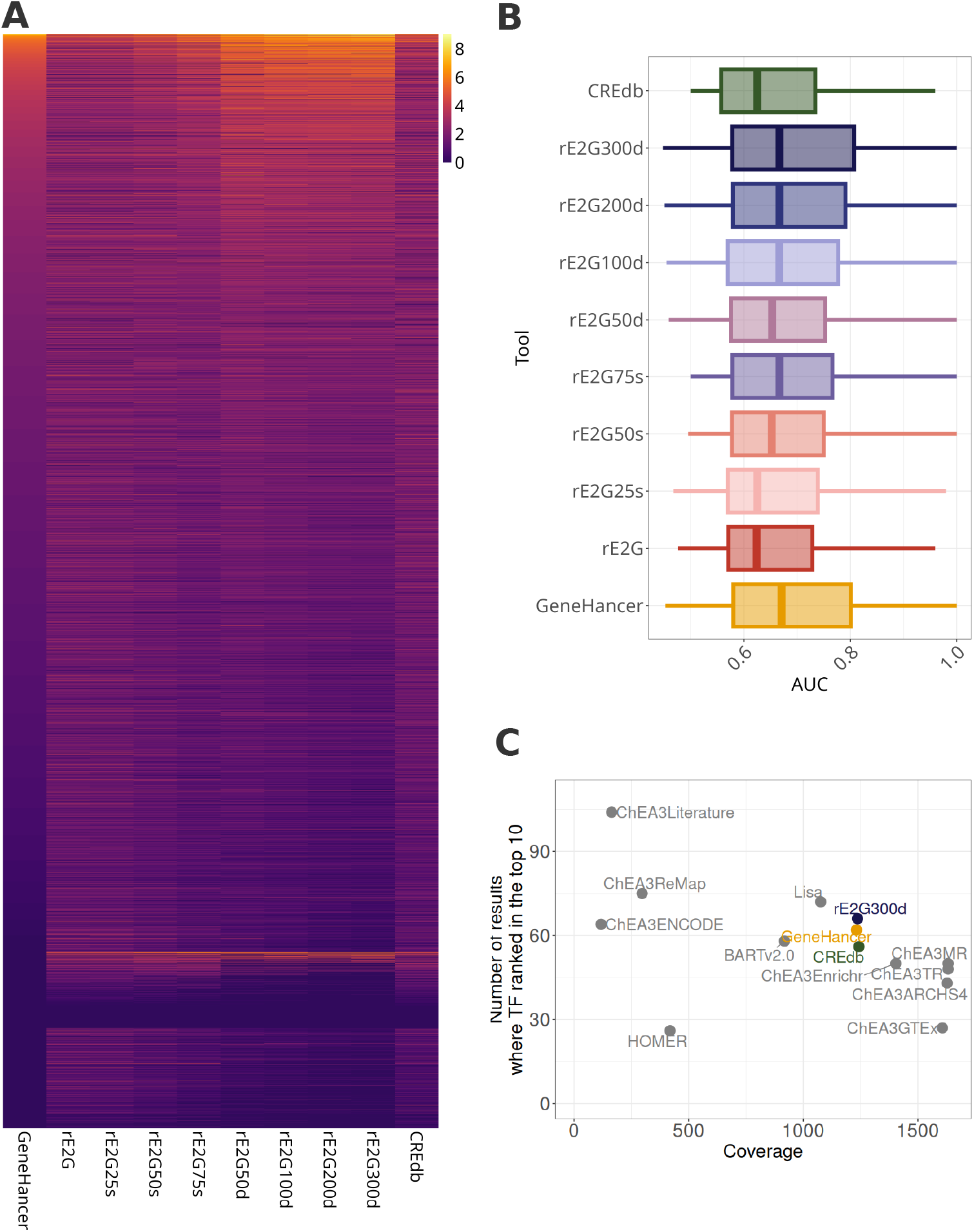
Performance evaluation of the new CRE–gene association databases. (A) Heatmap showing log_2_ odds ratio from TF ChIP-seq profiles corresponding to the TFs specified in the 369 benchmarking gene sets. Each row represents a ChIP-seq experiment for a TF associated to a benchmarking set, and each column corresponds to a different version of TFEA.ChIP. Non-significant enrichments (adjusted p *>*= 0.05) are shown in dark purple for visual clarity. Rows are ordered by enrichment strength in GeneHancer, from highest to lowest. (B) Boxplots summarizing the distribution of area under the curve (AUC) values across all benchmarking sets for each TFEA.ChIP version. AUC values were computed over 100 iterations per gene set, each using a different randomly sampled set of negative TF labels (see Methods). The distributions of mean AUC values across iterations are shown. (C) Comparative performance of TFEA.ChIP versus alternative TF enrichment tools. The x-axis indicates the coverage of each tool (i.e., the number of TFs analyzed), and the y-axis shows the number of benchmarking gene sets where the relevant TF ranked within the top 10 results.

We further benchmarked TFEA.ChIP against four widely used TF enrichment tools: BARTv2.0 [12], Lisa [15], ChEA3 and HOMER [7]. Using the 369 MSigDB gene sets as benchmarking [10, 18], we assessed each tool’s ability to identify the correct TF and rank it among the top 10 predictions. TFEA.ChIP demonstrated a strong balance between prediction performance and TF coverage, producing results for over 1,200 TFs (Figure 2C). These results confirm that the updated TFEA.ChIP framework not only surpasses its original version but also outperforms other leading tools. By generating stronger enrichment signals and more reliably identifying true regulatory TFs, it represents a robust and effective method for TF enrichment analysis.

### Case study: hypoxia meta-analysis

Hypoxia is a well-characterized cellular stress response driven primarily by hypoxia-inducible factors (HIFs), a family of TFs composed of a dimer between an alpha and a beta subunit. Among the alpha subunits, HIF1A and EPAS1 (also known as HIF2A) are the most studied and play central roles in regulating the transcriptional response to low oxygen conditions [2]. To evaluate the biological relevance and effectiveness of the updated TFEA.ChIP implementation, we applied it to a consensus hypoxia gene signature derived from a meta-analysis of 394 RNA-seq samples across 46 independent studies [14]. This meta-analysis estimated the pooled transcriptional effect of hypoxia in human cell lines cultured in vitro, generating a robust gene expression profile characteristic of the hypoxic response. We used this gene set to test whether TFEA.ChIP could correctly prioritize HIF-related TFs using two complementary enrichment approaches: ORA and GSEA.

In the ORA framework, the updated TFEA.ChIP databases, especially those filtered by depth, yielded significantly higher OR for HIF-related TFs compared to the original GeneHancer version for ChIP-seq datasets where the hypoxia pathway was known to be active (Figure 3A, 3B). Conversely, datasets where the pathway was inactive (e.g., ChIP-seq from untreated samples in normoxia) showed lower log_2_OR, suggesting improved specificity and reduced false positives. This indicates that the updated regulatory maps are more accurate in capturing biologically relevant TF–gene associations.

**Fig. 3.**
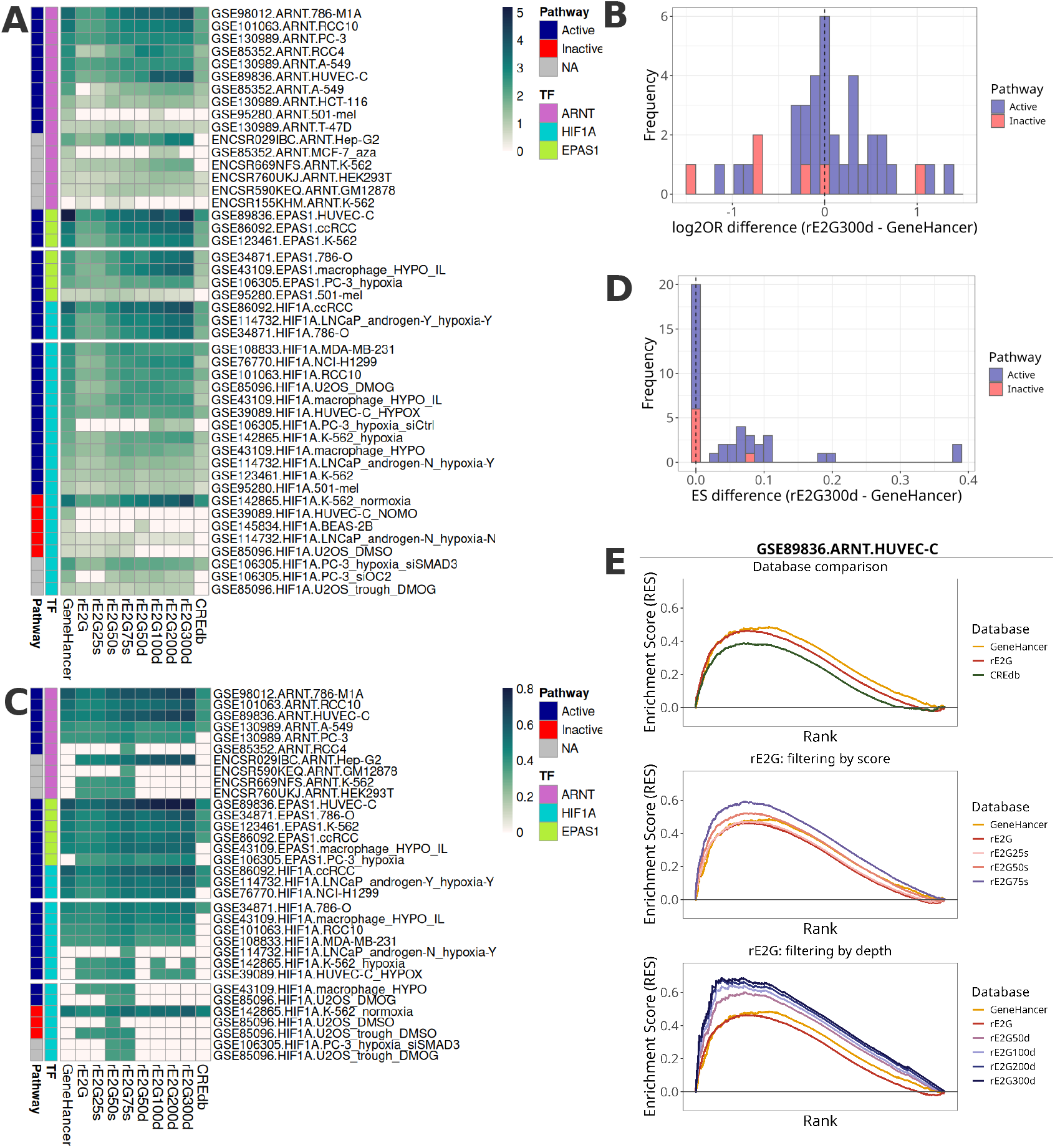
Evaluating TFEA.ChIP performance using a hypoxia meta-analysis signature. (A) Heatmap showing the log_2_ odds ratio from the Overrepresentation Analysis using the hypoxia gene signature derived from a meta-analysis. Only HIF ChIP-seq datasets that produced a statistically significant enrichment (adjusted p-value *<* 0.05) with at least one version of the databases are shown. Non-significant values (adjusted p *>*= 0.05) are shown in white. Rows are ordered first by TF, then by pathway, and finally by enrichment strength in GeneHancer, sorted from highest to lowest. Annotation bars indicate HIF pathway activation status (blue if “Active”, red if “Inactive”, grey if “Not assigned”) and the TF subunit (purple if “ARNT”, blue if “HIF1A”, green if “EPAS1”). (B) Histogram of the difference in log_2_OR between the best-performing rE2G database (depth *>* 300) and the original database. Bars are colored by HIF pathway status: blue for “Active”, red for “Inactive”. (C) Heatmap showing ES from GSEA using the same hypoxia meta-analysis signature. As in panel A, only HIF ChIP-seq datasets with at least one significant result are included. Non-significant enrichments are shown in white. Rows are ordered first by TF, then by pathway, and finally by enrichment strength in GeneHancer, sorted from highest to lowest. Pathway activation and TF subunit annotations are colored as described above. (D) Histogram of the difference in GSEA enrichment score between the best-performing rE2G version (depth *>* 300) and the original database, with bars separated by HIF pathway status. (E) Example of GSEA output using the GSE89836 ChIP-seq dataset for ARNT in HUVEC. The enrichment curves for the original GeneHancer database are shown in all plots for comparison. In the first plot, it is compared with CREdb and the unfiltered rE2G database. The second plot compares it with rE2G filtered by score, and the third with rE2G filtered by depth.

Given that the hypoxia meta-analysis provided a ranked gene list with effect sizes (i.e., weighted average LFC across studies), we also applied GSEA to assess enrichment. Again, the depth-filtered rE2G datasets outperformed both the original GeneHancer database and score-filtered versions, producing stronger and more consistent enrichment signals for HIF-related TFs in contexts where the pathway was active (Figure 3C, 3D). In contrast, the CREdb-based database yielded less compelling results, showing weaker enrichment for activated ChIP-seq datasets and failing to identify some as significantly enriched. This suggests limited sensitivity to pathway activation and decreased utility for fine-grained regulatory analysis. A representative example is the enrichment of ARNT using ChIP-seq data from HUVEC (Figure 3E). Although enrichment was significant across all methods, the depth-filtered rE2G versions outperformed others, yielding the strongest signals. This underscores their improved resolution and functional relevance in this biologically meaningful context.

These results show that the updated CRE–gene association strategies, particularly depth-based filtering of rE2G, substantially enhance TFEA.ChIP’s ability to identify key regulatory TFs from complex expression data. In the case of hypoxia, these improvements translate into more precise, sensitive, and interpretable enrichment outputs. This case study highlights the value of integrating experimentally validated and context-aware regulatory maps like ENCODE’s rE2G for advancing TF enrichment analysis.

## Discussion

TF enrichment analysis is a pivotal step in interpreting gene expression profiles and identifying upstream regulatory drivers. In this study, we present an enhanced version of TFEA.ChIP, incorporating CRE–gene association data from ENCODE’s rE2G predictive models [4]. This update introduces both methodological and functional improvements, including: the integration of recurrence-based regulatory maps, (2) robust benchmarking using curated gene sets, (3) improved performance relative to other tools, and (4) an expanded R/Bioconductor interface enabling seamless integration with standard differential gene expression analysis workflows.

A core challenge in TF enrichment analysis lies in accurately linking CREs to their target genes. These regulatory connections are often highly context-dependent, particularly for enhancers, which tend to be active in a tissue or cell type-specific manner. For example, only approximately 14% of H3K27ac-marked regions are shared across human tissues, while about 62% are tissue-specific [8]. Shared regulatory elements are typically promoter-proximal and linked to broadly expressed housekeeping genes, whereas tissue-specific elements tend to be distal enhancers that control specialized gene expression programs. Although the global chromatin landscape is relatively stable, enhancer activity and enhancer–gene interactions vary substantially between biological contexts. Then, in an ideal scenario, TF enrichment tools would use cell type-specific CRE–gene associations and ChIP-seq data to fully capture this regulatory complexity. However, such context-specific datasets remain limited and unevenly distributed across TFs and tissues, posing a significant challenge for enrichment tools. To address this limitation, TFEA.ChIP adopts a generalized CRE–gene linking strategy that integrates associations observed across multiple biosamples. Although this method does not capture detailed, context-specific regulation, it prioritizes robust and broadly conserved TF-gene interactions. As a result, TFEA.ChIP provides consistent and reliable TF enrichment results across diverse biological conditions, even without cell type-specific regulatory input. Unexpectedly, the generalized CRE-gene linking strategy outperformed cell-type specific approaches. For example, when testing TFEA.ChIP versions built with TF–gene associations tailored to MCF7 and A549 cells using ENCODE’s rE2G models, we observed weaker signals for the expected TFs compared to those obtained with the generalized model (Figure S2). This finding highlights the strength of a generalizable framework, especially considering that the ChIP-seq data used by TFEA.ChIP is derived from a diverse set of cell types rather than a single, context-specific source.

The original version of TFEA.ChIP was based on GeneHancer, a generalized CRE–gene association database that, like CREdb, aggregates regulatory links across multiple cell types. In contrast, the ENCODE rE2G predictive models offer cell type-specific CRE–gene associations. To adapt these cell type-specific models for broader use, we compiled a unified compendium of all available rE2G predictions and applied filtering strategies to generalize the data. Specifically, we filtered CRE–gene links based on functional evidence supporting the regulatory interaction and (ii) their recurrence across biosamples. This approach enabled us to transform the highly specific rE2G predictions into a robust, context-agnostic dataset better suited for general-purpose TF enrichment analysis. To evaluate the impact of these refinements, we benchmarked the updated version of TFEA.ChIP using a curated set of 369 gene sets as well as a focused case study on hypoxia-responsive genes. Notably, CREdb underperformed compared to the previous GeneHancer-based version, suggesting that including large numbers of highly cell type-specific links can actually dilute enrichment signals, potentially due to increased TF target coverage at the expense of specificity. The unfiltered rE2G compendium performed comparably to CREdb, but further improvements were achieved through filtering. While raising the confidence score threshold led to modest gains, recurrence-based filtering produced substantially better results. In particular, the rE2G dataset filtered to include only links present in at least 300 biosamples (rE2G300d) yielded the strongest enrichment signals and consistently outperformed both the original TFEA.ChIP implementation and all other filtering strategies. These findings suggest that recurrence across diverse biological contexts can serve as a useful proxy for regulatory relevance, particularly when both TF binding data and CRE–gene associations originate from heterogeneous sources. While the benefits of this approach are not uniformly observed across all contexts, it can enhance enrichment in specific cases. For example, in the hypoxia case study, it led to stronger enrichment signals for HIF subunits in conditions where the pathway was active, and absent signals when it was inactive. Finally, we compared the updated TFEA.ChIP against several widely used TF enrichment tools, including BARTv2.0, Lisa, ChEA3 and HOMER. TFEA.ChIP demonstrated an effective trade-off between TF coverage and performance. These results highlight the advantages of incorporating experimentally derived CRE–gene links over heuristic or distance-based models for robust and interpretable TF enrichment analysis.

All CRE–gene association resources evaluated in this study are derived from human genomic data. While this limits TFEA.ChIP to regulatory interactions mapped in humans, the tool nonetheless performed well on gene sets derived from mouse studies. This cross-species applicability likely reflects the evolutionary conservation of key transcriptional regulatory mechanisms. Indeed, multiple studies, including those from the ENCODE Consortium, have shown that many candidate enhancers in the mouse genome have orthologous counterparts in humans [19]. These human orthologs often retain similar chromatin signatures and regulatory functions [13], and in many cases, functional conservation persists even in the absence of sequence conservation [17], particularly when the elements occupy syntenic genomic regions. Nonetheless, certain limitations remain. The exclusive reliance on human-derived regulatory maps may reduce the tool’s precision when applied to non-human datasets, especially in the case of tissue-specific enhancers that lack conserved orthologs. Moreover, while recurrence-based filtering improves generalizability, it may overlook context-specific interactions that are biologically meaningful but rare. Future versions of the tool could incorporate context-aware or species-specific regulatory maps to better balance specificity, coverage, and cross-species applicability.

Functionally, the updated TFEA.ChIP now offers several new features that enhance usability and integration with downstream workflows. The R/Bioconductor version supports: direct input of differential expression results from widely used packages such as DESeq2, edgeR and limma, (2) a wrapper function for streamlined end-to-end analysis, (3) meta-analyses to aggregate results from multiple ChIP-seq datasets for the same TF, and (4) filtering of TFs not expressed in the input sample, reducing false positives and increasing interpretability. These additions make TFEA.ChIP more accessible and adaptable for routine transcriptomic analyses.

In summary, this updated version of TFEA.ChIP offers a robust, biologically grounded, and scalable framework for TF enrichment analysis. By integrating high-confidence and recurrent CRE–gene associations, validating its performance across diverse benchmarks, and expanding user-facing functionality, TFEA.ChIP provides a valuable and practical resource for decoding transcriptional regulation from gene expression data.

## Conclusion

The updated TFEA.ChIP introduces enhanced CRE–gene association strategies that improve the accuracy and generalizability of TF enrichment analysis. Recurrence-based filtering of ENCODE rE2G data, in particular, produced the most robust results, outperforming both the original implementation and alternative tools. TFEA.ChIP’s R-based implementation enables easy integration into differential expression workflows, providing a practical and scalable tool for transcriptomic regulatory analysis. These advances establish TFEA.ChIP as a reliable and accessible tool for decoding transcriptional regulation across diverse biological contexts.

## Author contributions statement

Y.B. implemented the updates to TFEA.ChIP, performed all analyses, and wrote the manuscript. Y.B. and L.P. conceived the study, developed the methodology, interpreted the results and revised the manuscript. Both authors read and approved the final version.

## Acknowledgments

We thank Dr. Wyeth Wasserman for his thoughtful input and stimulating discussions on the regulatory genomics concepts underlying this work.

## Data availability

The TFEA.ChIP package is available through the Bioconductor repository. The development version of the package, including source code, documentation, and example workflows, is openly accessible on GitHub at https://github.com/yberda/TFEA.ChIP. An interactive web-based version is also provided as a Shiny application at https://www.iib.uam.es/TFEA.ChIP/, allowing users to perform TF enrichment analysis without requiring local installation.

