## Supplementary Figures for "Enhancing TFEA.ChIP with ENCODE Regulatory Maps for Generalizable Transcription Factor Enrichment"

### Supplementary Material

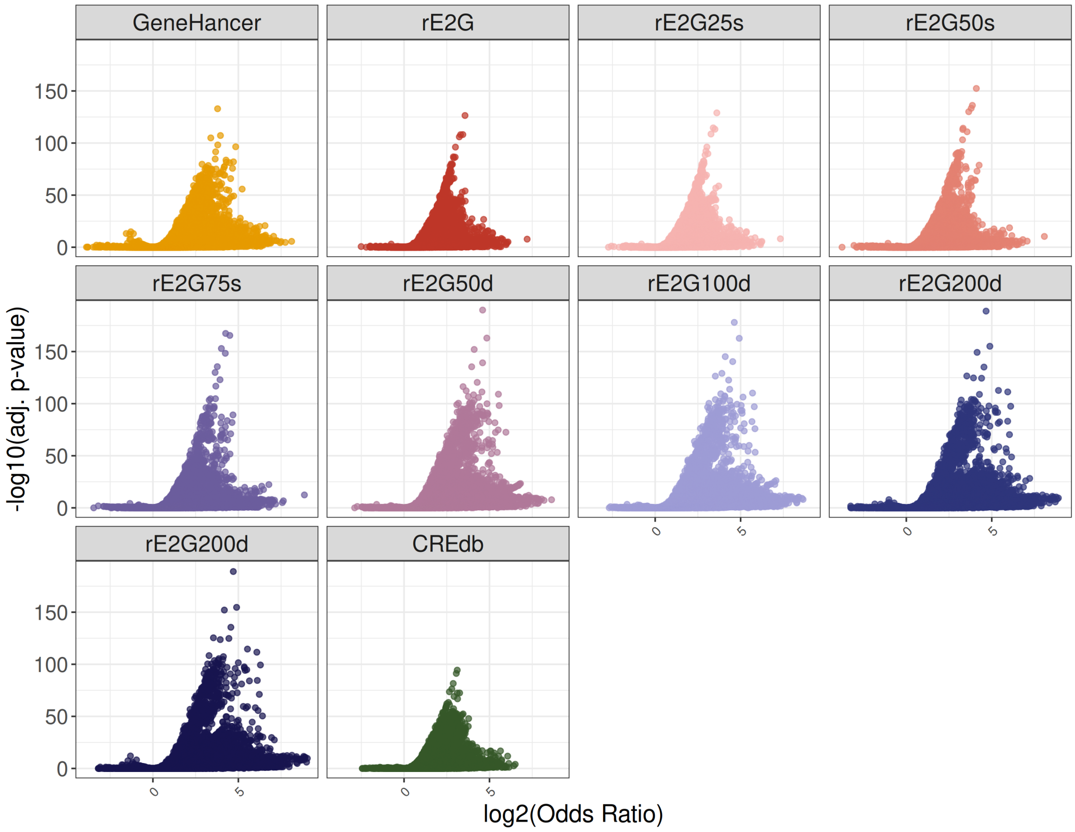


Figure S1: Volcano plots summarizing TF enrichment results across different CRE–gene association databases in TFEA.ChIP. Each panel shows enrichment results for one database: the original GeneHancer and multiple updated versions of the rE2G models and CREdb. Each point represents the enrichment outcome for a TF from a ChIP-seq dataset that is expected to be enriched, based on a known benchmarking gene set. The x-axis shows the $\log_2$ odds ratio indicating the strength of enrichment, and the y-axis shows the -$\log_2$(adjusted p-value), showing statistical significance.

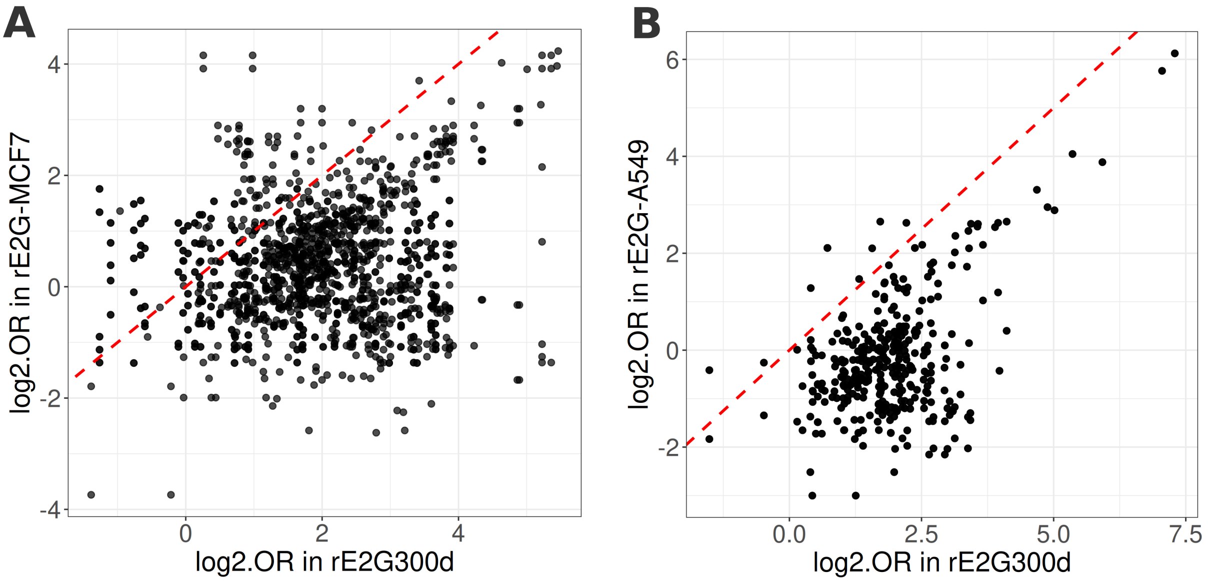


Figure S2: Comparison of TF enrichment using general versus cell type-specific rE2G models in TFEA.ChIP. $Log_2$ odds ratio values from Overrepresentation Analysis are shown for TF ChIP-seq datasets expected to be enriched based on benchmarking gene sets. Panel A uses MCF7-derived gene sets, and Panel B uses A549-derived sets. The red dashed line indicates the identity line ($y = x$), marking equal enrichment by the general (rE2G300d) and cell type-specific models.
